# Experimental evolution of a reduced bacterial chemotaxis network

**DOI:** 10.1101/2024.03.14.584839

**Authors:** Manika Kargeti, Irina Kalita, Sarah Hoch, Maryia Ratnikava, Wenhao Xu, Bin Ni, Ron Leonard Dy, Remy Colin, Victor Sourjik

## Abstract

Chemotaxis allows bacteria to follow chemical gradients by comparing their environment over time and adjusting their swimming behavior accordingly. The chemotaxis signaling pathway is highly conserved among all chemotactic bacteria. The system comprises two modules: one for environmental sensing and signal transduction toward the flagellar motor, and the other for adapting to the constant level of background stimulation and providing short-term memory for temporal comparisons. Previous experimental analysis and mathematical modeling have suggested that all components of the paradigmatic chemotaxis pathways in *Escherichia coli* are essential. This indicates that it may contain a minimal set of protein components necessary to mediate gradient sensing and behavioral response. To test this assumption, here we subjected strains carrying deletions in chemotaxis genes to experimental laboratory evolution. We observed that the core components of the chemotaxis pathway are indeed essential. However, the absence of individual auxiliary pathway proteins, including the adaptation enzymes that are conserved in a vast majority of bacteria, and the phosphatase, could be compensated for to varying degrees by changes in other pathway components. Our results suggest that the experimental evolution of these deletion strains has led to the emergence of alternative strategies for bacterial chemotaxis, demonstrating the surprisingly rapid evolvability of this signaling network.

## Introduction

Most motile bacteria can follow gradients of nutrients and other stimuli in their environment through chemotaxis. This process is crucial for bacterial physiology, including growth optimization, collective behaviors, and interactions with eukaryotic hosts (1, 2). The central part of the signaling pathway responsible for bacterial chemotaxis is highly conserved among prokaryotes (3, 4). *Escherichia coli* has one of the simplest chemotaxis pathways, consisting almost exclusively of highly conserved proteins, and it is one of the most studied quantitative models for signal transduction in biology (5). The mechanism of bacterial chemotaxis relies on temporal comparisons of swimming bacteria, where based on the perceived changes in environmental conditions, the chemotaxis signaling system determines whether the bacterium should continue running in its current direction or reorient itself (6). This strategy is thought to require two modules: one for rapid environmental sensing and signal transduction, and another for slower adaptation that enables short-termed temporal comparisons of environmental conditions (5, 7).

The environmental sensory module (Figure 1A) comprises transmembrane receptors, also known as methyl-accepting chemotaxis proteins, that control the autophosphorylation activity of the receptor-associated kinase CheA with the assistance of the scaffolding protein CheW (8). Out of five *E. coli* chemoreceptors, at least one of the two major transmembrane chemoreceptors, Tar or Tsr, is required for the proper formation of sensory complexes and regulation of CheA. The sensory module’s output is transmitted to the flagellar motor through the CheA-dependent phosphorylation of the response regulator CheY. In *E. coli*, the phosphorylation of CheY and its binding to flagellar motors increase when the bacterium travels in an unfavorable direction. This induces a switch in the motor rotation from the default counterclockwise (CCW) to clockwise (CW) direction, resulting in the flagellar bundle falling apart and the bacterium tumbling and reorientating. When swimming in a favorable direction, such as traveling up the gradient of attractant, the binding of attractant to receptors inhibits CheA autophosphorylation, which reduces CheY phosphorylation and, in turn, favors CCW rotation and smooth swimming. This core of the sensory and signaling module is conserved in all bacterial chemotaxis systems and is evolutionary related to the broader class of bacterial two-component pathways (9). In addition, the chemotaxis signaling module of *E. coli* and closely related proteobacteria includes the phosphatase CheZ that is responsible for the rapid dephosphorylation of CheY, whereas other chemotaxis systems contain alternative phosphatases.

**Figure 1.**
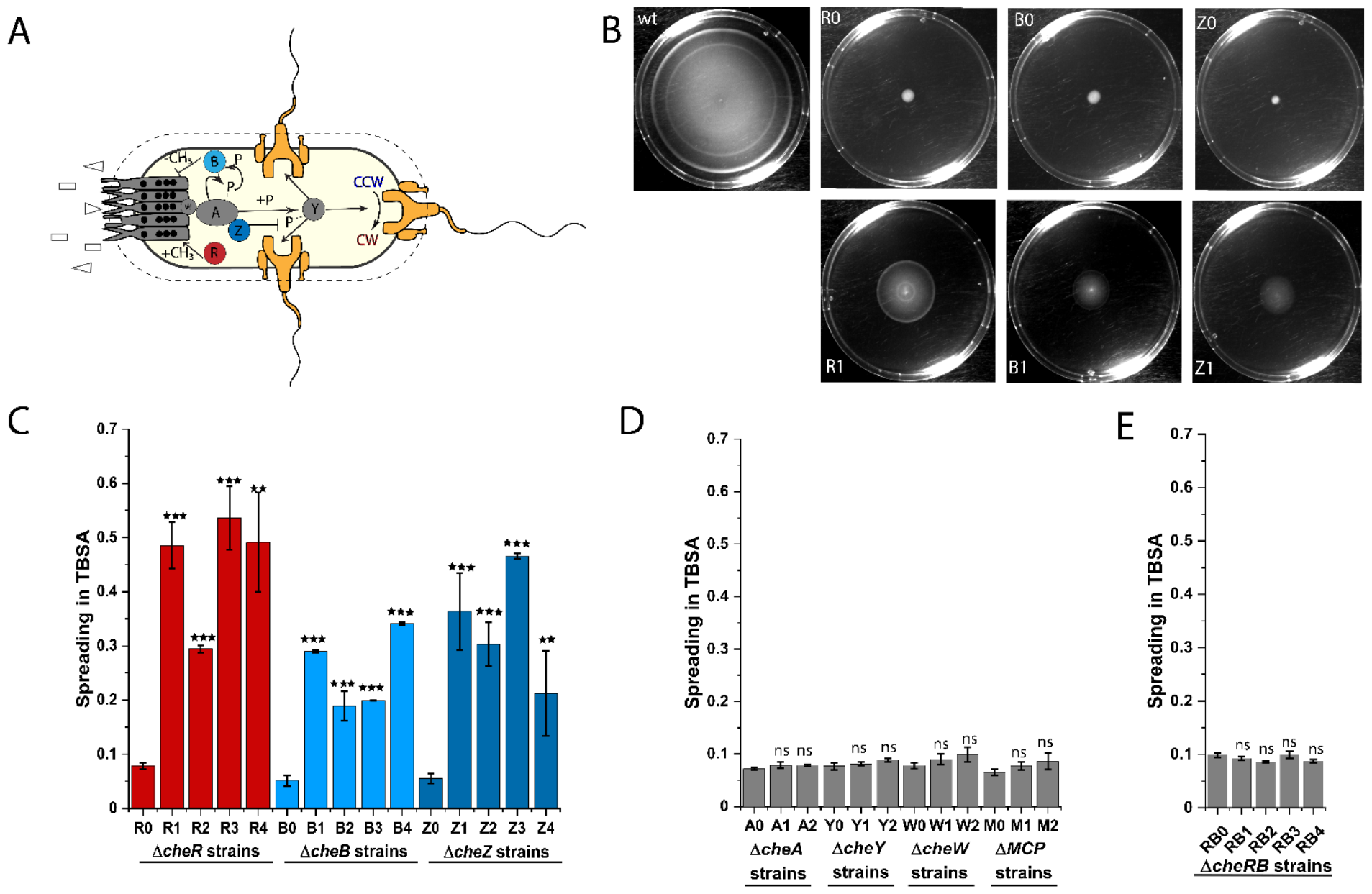
Experimental evolution partly restores spreading of chemotaxis mutants in soft agar. **(A)** Schematic representation of the chemotaxis signaling pathway of *Escherichia coli*. The pathway includes transmembrane chemoreceptors (two major chemoreceptors, Tar and Tsr, are shown) that form sensory complexes together with a kinase CheA (A) and an adaptor CheW (W). CheY (Y) phosphorylation by CheA mediates signal transduction to flagellar motor, inducing a switch from the default counterclockwise (CCW) to clockwise (CW) rotation. The adaptation module, including CheR (R) and CheB (B), regulates chemoreceptor activity through methylation and demethylation of chemoreceptors on four specific glutamates (black circles). The phosphatase CheZ (Z) is responsible for rapid CheY dephosphorylation. See text for more details. **(B)** Spreading of the wildtype (wt) *E. coli* RP437, of its Δ*cheR* (R), Δ*cheB* (B) or Δ*cheZ* (Z) derivatives deleted for individual chemotaxis genes, either before (denoted as “0”) or after evolution for 30 days in TB soft agar (TBSA), with an example of one evolved line (denoted as “1”) shown for each strain. **(C-E)** Size of the spreading colonies for Δ*cheR* (R), Δ*cheB* (B) and Δ*cheZ* (Z) strains (C); for Δ*cheA* (A), Δ*cheW* (W) and Δ*mcp* (M) strains (D); and for Δ*cheR cheB* (RB) strain (E) normalized to that of the wildtype *E. coli* RP437. Spreading was measured for three independent replicates after incubation for ∼16 h in TBSA. Several independent lines of evolution (indicated by numbers) are shown for each strain. Error bars indicate standard errors. *P* values were calculated for comparisons between spreading of evolved and respective non-evolved strains for each deletion, using two tailed t-test (ns, not significant; ^*^, *P <* 0.05; ^**^, *P <* 0.01; ^***^, *P <* 0.001).

The adaptation module comprises two enzymes, the methyltransferase CheR and the methylesterase CheB, which respectively methylate or demethylate four specific glutamates on chemoreceptors. Unmethylated glutamates promote a low activity state, whereas methylated glutamates promote a high activity state of chemoreceptors. The receptors are first expressed in the intermediate activity state, with two of the four methylation sites being encoded as glutamines, which funciton similarly to methylated glutamates and are deamidated by CheB to glutamates. This adaptation module is unique among bacterial two-component signaling pathways but it is nearly universally present in chemotaxis systems, with the notable exception of gastric species of *Helicobacter* (10). Enzymatic activity of the methylation enzymes depends on the receptor activity state, and the resulting negative feedback ensures that the steady-state activity of the pathway can adapt to intermediate level even in the presence of persistent stimulation. Additionally, changes in methylation occur with a delay following receptor stimulation, creating a short-term memory that swimming bacteria use for temporal comparisons of environmental conditions. Both functions of the adaptation module are assumed to be essential for efficient bacterial chemotaxis.

Efficient chemotaxis in *E. coli* requires all cytoplasmic chemotaxis proteins and at least one major chemoreceptor (11). One reason for that is the extreme tumbling bias observed in cells lacking these either of these proteins. Strains with deletions in *cheW, cheA, cheY*, or all receptor genes do not phosphorylate CheY, resulting in continuous running without reorientations. Conversely, *cheZ*-deficient cells have an excess of phosphorylated CheY (CheY-P) and tumble most of the time. Deletions in *cheR* or *cheB* genes result in very low or high levels of pathway activity, and thus of tumbling bias, and disabling adaptation and temporal comparisons by the chemotactic cells. As a result, these mutants are unable to efficiently navigate chemical gradients in liquid (12). They also have a deficiency in spreading on soft agar plates (13), which is a commonly used assay for motility and chemotaxis that relies on the spreading of motile bacteria through agar pores following self-generated gradients of consumed chemoattractant nutrients (14).

Although *cheR* and *cheB* mutants appear to be unable to perform chemotaxis, early studies indicated that *E. coli* strains lacking both CheR and CheB activities may exhibit some degree of tactic behavior (13, 15). This was further supported by the emergence of spontaneous (pseudo)revertants of the *cheR* deletion strain that could spread on soft-agar plates, with compensatory mutations mapping to either *cheB* (16) or *tsr* (17) genes. However, the compensatory mechanisms underlying this phenomenon remained unclear (17), and a subsequent study concluded that the *cheR cheB* mutants or the *cheR* revertants may rather spread in soft agar in a chemotaxis-independent fashion due to their intermediate tumbling bias, which enables slow and non-directional movement through the agar pores (14). Such pseudotactic mutants, which carry mutations in genes encoding flagellar hook or motor proteins, have also been isolated in chemotaxis-deficient strains of other bacteria (18).

To investigate whether all *E. coli* pathway proteins, including adaptation enzymes, are indeed essential for chemotaxis, we experimentally evolved a set of *E. coli* deletion strains for chemotaxis-driven spreading in soft agar over several hundred generations. Experimental evolution, also known as adaptive laboratory evolution, is a powerful approach to investigate how individual proteins and gene regulatory networks adapt under defined selection pressure (19, 20). Recently, it has been used to investigate the evolvability of genetic regulation under selection for motility (21) and the underlying cost-benefit tradeoffs between motility and growth (22-24). Here we tested the capability of compensating the absence of individual proteins by evolutionary remodeling the chemotaxis pathway. Our results demonstrate that the absence of auxiliary chemotaxis proteins could be reproducibly compensated by short-term adaptive evolution, albeit to varying extents, while core signaling functions remain essential. Importantly, the evolved *cheR* strain not only regained the ability to spread in soft agar but also demonstrated biased drift in chemical gradients in liquid, indicating its capability for true chemotaxis. This suggests that not only individual enzymatic activities and gene regulation, but also complex signaling pathways in bacteria are highly evolvable, allowing for the emergence of novel simplified chemotaxis strategies that bypass the lack of normally essential individual components.

## Results

### Experimental evolution can compensate for defects caused by the deletions of several chemotaxis genes

Experimental evolution of *E. coli* mutant strains was performed under selection for increased spreading on tryptone broth soft-agar (TBSA) plates for 30 cycles of up to 16 hours each (Figure S1A), and with up to four independently evolved lines. We observed that the spreading of the evolved Δ*cheR*, Δ*cheB*, and Δ*cheZ* strains, which lack either the individual adaptation enzymes or the phosphatase (Figure 1A), largely improved compared to that of the original deletion strains (Figure 1B,C). In contrast, the absence of core components of the chemotaxis pathway, including CheA, CheY, CheW, or all chemotaxis receptors, could not be compensated for by short-term evolution (Figure 1D and Figure S1B). Furthermore, no improvement in the spreading of a double Δ(*cheR cheB*) deletion strain under selection was observed (Figure 1E). However, the spreading of this deletion strain was slightly better than that of the individual *cheR* or *cheB* deletion strains, which is consistent with the previous report (14). As the evolved Δ*cheR* strain showed the largest enhancement of spreading, we evolved four additional Δ*cheR* lines, all of which showed a similar enhancement in spreading (Figure S2).

### Evolved strains exhibit compensatory changes in motility

We next investigated changes in the motility phenotypes of the evolved *E. coli* strains, by tracking cell swimming in liquid. Consistent with previous finding on the importance of intermediate tumbling frequency for spreading in soft agar (14), the evolved strains exhibited compensation for the defects in tumbling that were present in the original deletion strains, so that the fraction of time that cells spent tumbling became more similar to the wildtype (Figure 2A). All four evolved Δ*cheR* lines became more tumbly than the original (R0) Δ*cheR* strain, whereas all four evolved Δ*cheZ* and two Δ*cheB* lines became less tumbly. This modulation of tumble bias by experimental evolution correlated well with increased spreading in TBSA (Figure 2B), suggesting that the wildtype tumble bias is optimal for spreading.

**Figure 2.**
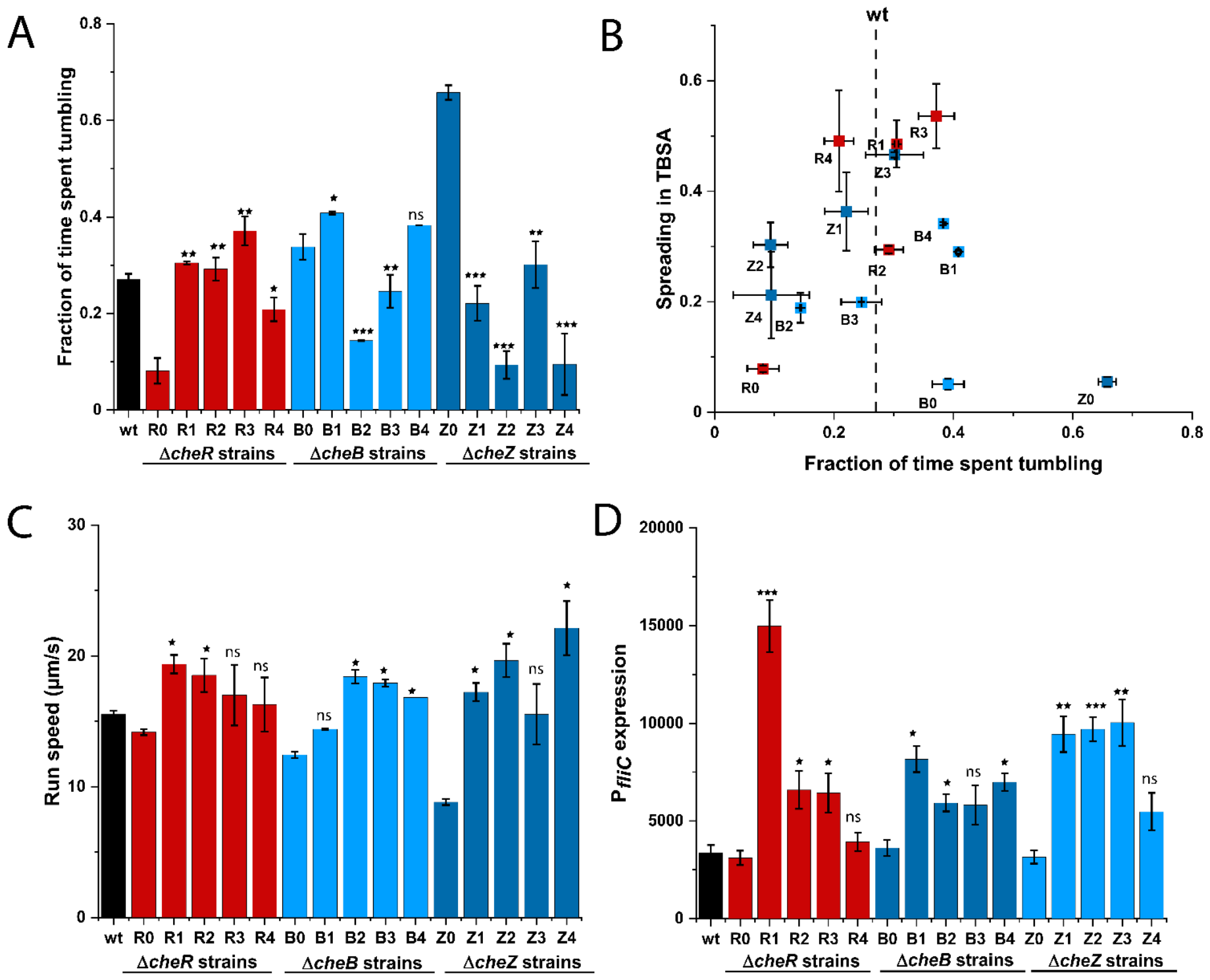
Evolved changes in motility phenotypes and flagellar gene expression. **(A)** Fraction of time spent tumbling for wt, parental strains (R0, B0 and Z0), and evolved strains (R1-R4, B1-B4, Z1-Z4), measured in three independent replicates. Significance analysis was done in comparison to respective non-evolved strains. **(B)** Spreading in TBSA (data from Figure 1C) plotted as a function of the fraction of time spent tumbling for individual strains. The dotted line indicates fraction of time spent tumbling for wt. **(C)** Run speed between two consecutive tumbles, measured for all strains in three independent replicates. Significance analysis was done in comparison to wt. Motility phenotypes were assessed using cell tracking (see Methods). **(D)** Activity of transcriptional *fliC* promoter (P*fliC*) reporter, measured as fluorescence of green fluorescent protein (GFP) using flow cytometry in three independent replicates. Significance analysis was done in comparison to respective non-evolved strains. Error bars indicate standard errors. *P* values were calculated using two tailed t-test (ns, not significant; ^*^, *P <* 0.05; ^**^, *P <* 0.01; ^***^, *P <* 0.001).

Furthermore, nearly all of the evolved strains exhibited a higher swimming velocity than the wildtype strain (Figure 2C). This increase in velocity was previously observed during the evolution of wildtype cells for spreading in TBSA, as a consequence of the elevated expression of the flagellin gene *fliC* and other flagellar genes (22). Therefore, we measured the activity of the transcriptional *fliC* promoter (P*fliC*) reporter, which reflects the expression of flagellar genes (22). The activity of this reporter was significantly higher in most of the evolved strains (Figure 2D), confirming that their increased swimming velocity is likely due to changes in flagellar gene expression.

### Specific compensatory mutations are observed in evolved strains

Whole genome sequencing revealed multiple mutations in the evolved Δ*cheR* (Table S1), Δ*cheB* (Table S2) and Δ*cheZ* (Table S3) lines. These mutations exhibited clear gene-deletion specific patterns and primarily affected chemotaxis or motility genes. The most prominent set of mutations in the evolved Δ*cheR* lines mapped to the *tsr* gene that encodes the serine-specific major chemoreceptor (Figure 3A and Table S1). Mutations in *tsr* were observed in seven out of eight lines. One line (R7) carried instead a mutation in the *tar* gene, which encodes the aspartate-specific receptor, and one line (R3) carried mutations in both *tsr* and *tar* (Figure 3B). The corresponding amino acid substitutions mapped to different receptor domains, including the ligand-binding domain, transmembrane helix, HAMP domain, methylation region, and in the signaling domain, similar to the previous report (17). These mutations may promote the active state of receptors, contributing to the suppression of the low-activity *cheR* phenotype. Indeed, M249 and L263 were previously shown to be critical for the helical packing of the HAMP bundle and for signal transduction (25), and T305 is located adjacent to the methylation site (26). Mutations in *tsr* may have been preferentially selected over those in *tar* because the first gradient followed by chemotactic cells on TBSA plates is that of the Tsr ligand serine (27), or because cells have higher levels of Tsr compared to Tar under our growth conditions.

**Figure 3.**
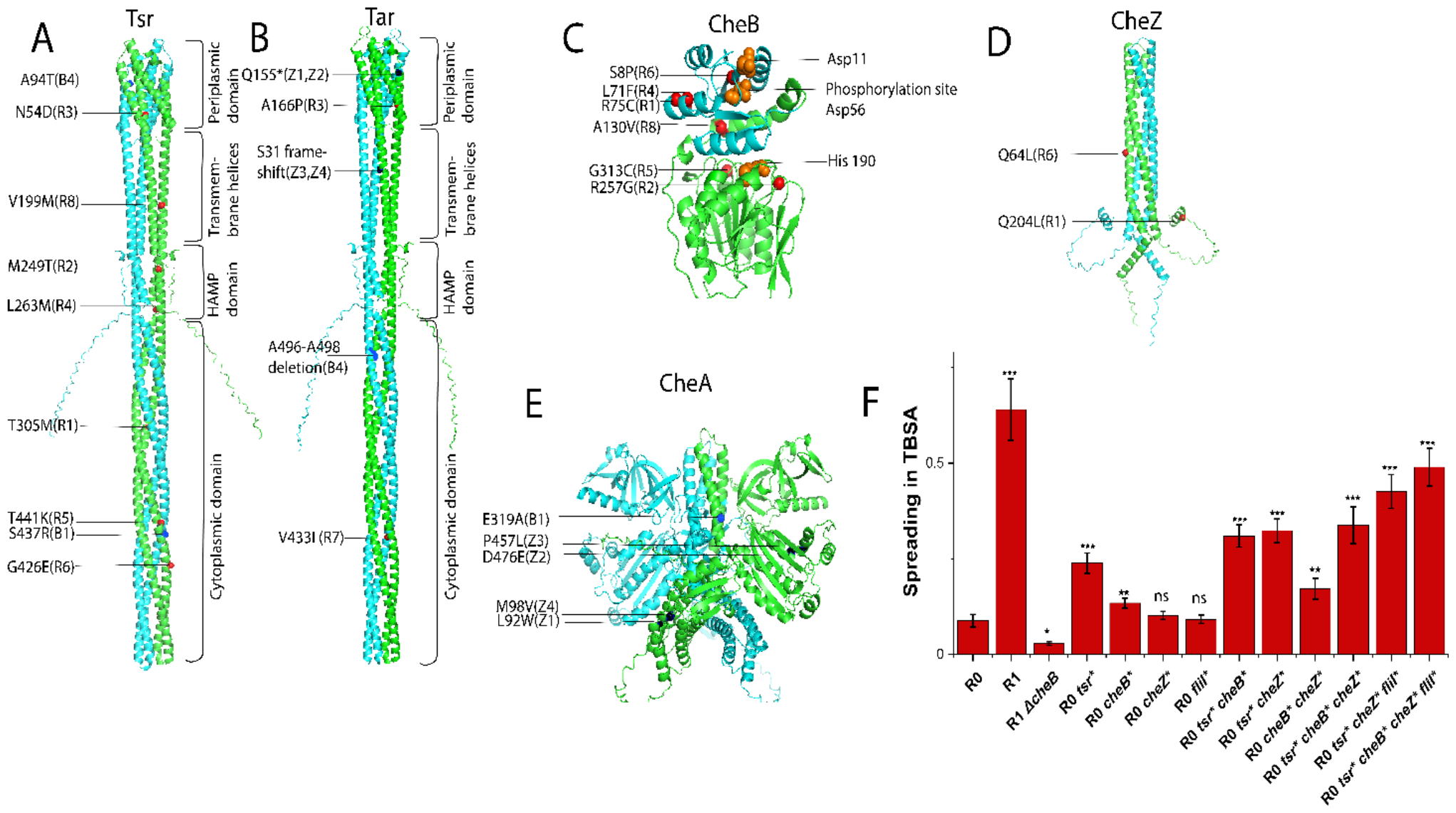
Evolutionary selected amino acid substitutions in chemotaxis proteins. Substitutions identified in evolved R strains (red), B strains (dark blue) and Z strains (black) in **(A)** Tsr, **(B)** Tar, **(C)** CheB, **(D)** CheZ, and **(E)** CheA, mapped on the respective protein structure. For chemoreceptors, functional domains are labeled. For CheB, residues in the phosphorylation pocket and in the catalytic pocket are marked in orange. **(F)** Spreading in TBSA of Δ*cheR* (R0) carrying individual mutations that were identified in the R1 strain, and their combinations, compared to spreading of R0 and R1 strains and of R1 strain carrying deletion of *cheB*. Values were measured in three independent replicates and normalized to spreading of wt. Error bars indicate standard errors. Significance analysis was in comparison to R0. *P* values were calculated using two tailed t-test (ns, not significant; ^*^, *P <* 0.05; ^**^, *P <* 0.01; ^***^, *P <* 0.001).

Most of the evolved Δ*cheR* lines also had mutations in the *cheB* gene (Figure 3C and Table S2). CheB consists of a regulatory CheY-like receiver domain, which is phosphorylated by CheA, and the enzymatic methylesterase domain (28). Mutations were found in both domains, with an apparent clustering near the catalytic pocket, the phosphorylation pocket, and the regulatory interface (29). These mutations may lower the methylesterase activity of CheB, as previously suggested (16), which could potentially compensate for the absence of methyltransferase activity. Nevertheless, no amber or frame-shift mutations were detected, indicating that some level of CheB activity is necessary for the restored spreading of the evolved Δ*cheR* lines.

Finally, two of the Δ*cheR* lines had mutations in *cheZ* (Figure 3D). Q204L affects the C-terminal CheY-binding peptide (30) and Q64L is close to the catalytic site of CheZ (31). Substitutions at both sites were reported to lower the phosphatase activity of CheZ (32), suggesting that they can partly offset the low kinase activity in the Δ*cheR* strain and thus increase the tumbling bias in the evolved lines.

Mutations in different domains of the receptor genes *tsr, tar* and *tap* were also found in all Δ*cheB* lines (Figure 3A,B and Table S2). Additionally, amino acid substitution were introduced in the dimerization domain of CheA and a short ALGD amino acid sequence was inserted in CheW (Figure 3E and Table S2). These mutations may affect the activity or stability of the ternary receptor-CheA-CheW complex, which could offset the hyperactive receptor phenotype caused by the Δ*cheB* deletion.

The evolved Δ*cheZ* lines compensated the hyperactive pathway phenotype, too, but acquired a different set of mutations (Table S3). In all lines, Tar translation was interrupted either by a stop codon mutation within the ligand-binding domain or by a frameshift in the transmembrane helix of the receptor (Figure 3B). Interestingly, each of these mutations was apparently acquired independently by two of the evolution lines. All Δ*cheZ* lines also had mutations in *cheA*, either in the P1 (phosphorylation site) domain or in the P4 (ATP binding kinase) catalytic domain (Figure 3E).

In addition to these mutations in the chemotaxis genes, almost all Δ*cheR* lines and one Δ*cheB* line had mutations in the genes that encode the export apparatus (*fliI*) and the basal body (*fliF, fliM, fliN* and *fliG*) of the flagellar motor (Table S1 and Table S2). Mutations in these genes have previously been shown to upregulate the expression of flagellar genes by enhancing secretion of the negative regulator (anti-sigma factor) FlgM (22). This may be consistent with the elevated *fliC* reporter activity in the evolved lines (Figure 2D). However, FliM, FliN and FliG have a dual role both in the function of flagella export apparatus and in the control of flagellar motor rotation, and could thus also directly affect cell tumbling. Evolved lines without mutations in the export apparatus or or the basal body gene also showed increased flagellar gene expression, possibly due to mutations or insertions in other genes such as *clpX, sspA* or *rpoD*, which are known to affect the expression of the flagellar regulon (22, 33) (Table S1, Table S2 and Table S3). Other observed mutations in *atp* genes encoding the PMF-dependent ATP synthase may increase motility by elevating the proton motive force.

We further investigated the order in which emergent mutations could be detected over the course of evolution for several Δ*cheR* lines (Figure S3). We observed that in all cases, the mutations in *tsr* were selected first, followed by mutations in other chemotaxis and/or flagellar genes.

We selected one of the best-spreading lines, R1, to evaluate the phenotypic impacts of individual mutations and their potential epistatic interactions. R1 carries mutations in the chemotaxis genes *tsr, cheB* and *cheZ* (Figure 3A,C,D), and a mutation in the flagellar export gene *fliI*. When these mutations were introduced individually into the Δ*cheR* strain, mutations in *tsr* or *cheB* significantly increased spreading on TBSA plates (Figure 3F). This result is consistent with these two genes being most commonly affected in the evolved Δ*cheR* lines. Spreading was further increased by combinations of multiple chemotaxis and *fliI* mutations. Interestingly, the observed order of selection for individual mutations during evolution of the R1 line (Figure S3A) is apparently consistent with the path of largest stepwise increase in spreading due to addition of each subsequent mutation, from *tsr* to *tsr cheZ* to *tsr cheZ fliI* to *tsr cheB cheZ fliI*. This spreading of the latter Δ*cheR* strain carrying four mutations in TBSA largely recapitulated that of the R1 line, with the residual difference being likely explained by an additional mutation in *sspA* gene present in this evolved strain, which was previously shown to increase flagellar gene expression (22).

We also investigated the effects of R1-specific mutations on the chemotactic spreading when introduced individually in the wildtype cells (Figure S3E). The introduction of *fliI* mutation, which is expected to increase expression of flagellar genes, led to enhanced spreading, consistent with the previous report (22). All other mutations in the wildtype background resulted in no or only modest changes in spreading, suggesting that all of the mutated genes remain well functional, including *cheB* and *cheZ* genes where mutations are expected to reduce the enzymatic activities of respective gene products.

In order to further directly test whether the function of CheB was required for the spreading of the R1 line, we introduced the *cheB* deletion in the evolved strain. Indeed, the R1 strain lacking *cheB* showed even less spreading in soft agar than the Δ*cheR* strain (Figure 3F), confirming our assumption that at least residual activity of CheB is necessary for the re-evolved spreading of Δ*cheR* strains.

Gradual increase in spreading was also observed when individual *cheB* and *tsr* mutations from R4 and R5 lines were introduced individually into the Δ*cheR* strain (Figure S3F). Similar to the R1-specific mutations, the effect of *tsr* mutations on spreading was stronger than that of *cheB* mutations, consistent with the tsr mutations being first to emerge in the populations of all tested evolved Δ*cheR* lines (Figure S3A-D).

### Evolved strains exhibit directional spreading in chemical gradients

Although previous studies concluded that spreading on TBSA plates does not necessarily require chemotaxis but only an intermediate cell tumbling frequency, such pseudotaxis is significantly slower than the chemotaxis-driven spreading (14, 34, 35). Given that the spreading of some of the evolved strains, in particular several Δ*cheR* lines, was highly efficient, even showing a clear ring at the edge of the spreading colony that is normally characteristic of chemotactic behavior (34, 35), we next investigated whether these strains might have regained the ability to perform chemotaxis. For that, we first used the M9 minimal medium soft agar (M9SA) gradient plates, containing glycerol as a carbon source, where the gradient was pre-established by diffusion after applying chemoeffector along the center line of the square plate (36, 37). On M9SA plates with the gradient of serine, the Tsr-specific attractant that is followed by the outer spreading ring in TBSA plates (27). Three out of four tested evolved Δ*cheR* lines showed biased spreading up the gradient similar to that of the wildtype strain (Figure S4A). This bias was confirmed by quantifying the ratio between the spreading distance toward the source and the distance away from the source (Figure S4D). Two out of four Δ*cheZ* lines also showed biased spreading up the gradient of serine, although less efficiently than that of the Δ*cheR* lines (Figure S4C and S4D). Other Δ*cheR* and Δ*cheZ* lines, as well as all Δ*cheB* lines (Figure S4B and S4D) showed little spreading or growth on M9SA plates, so their bias could not be determined.

Although the observed biased spreading up the serine gradient is indicative of chemotaxis, its interpretation is complicated by the fact that serine is metabolized, which could introduce growth bias on these gradient plates. We thus tested spreading of several Δ*cheR* lines using M9SA plates with a gradient of α-methyl-D, L-aspartate (MeAsp), a non-metabolizable analogue of aspartate. Indeed, a reproducible biased movement up the gradient of MeAsp could be observed for the R1, R4, and R5 lines, but not for R0 or the R3 line (Figure 4A and 4B). Notably, the R3 line carries an amino acid substitution (A166P) in the ligand binding domain of Tar (Figure 3B and Table S1), which might render it unable to sense MeAsp. Similarly, all Δ*cheZ* lines possess only a truncated version of Tar (Figure 3B and Table S3), and therefore could not be tested on MeAsp gradient plates.

**Figure 4.**
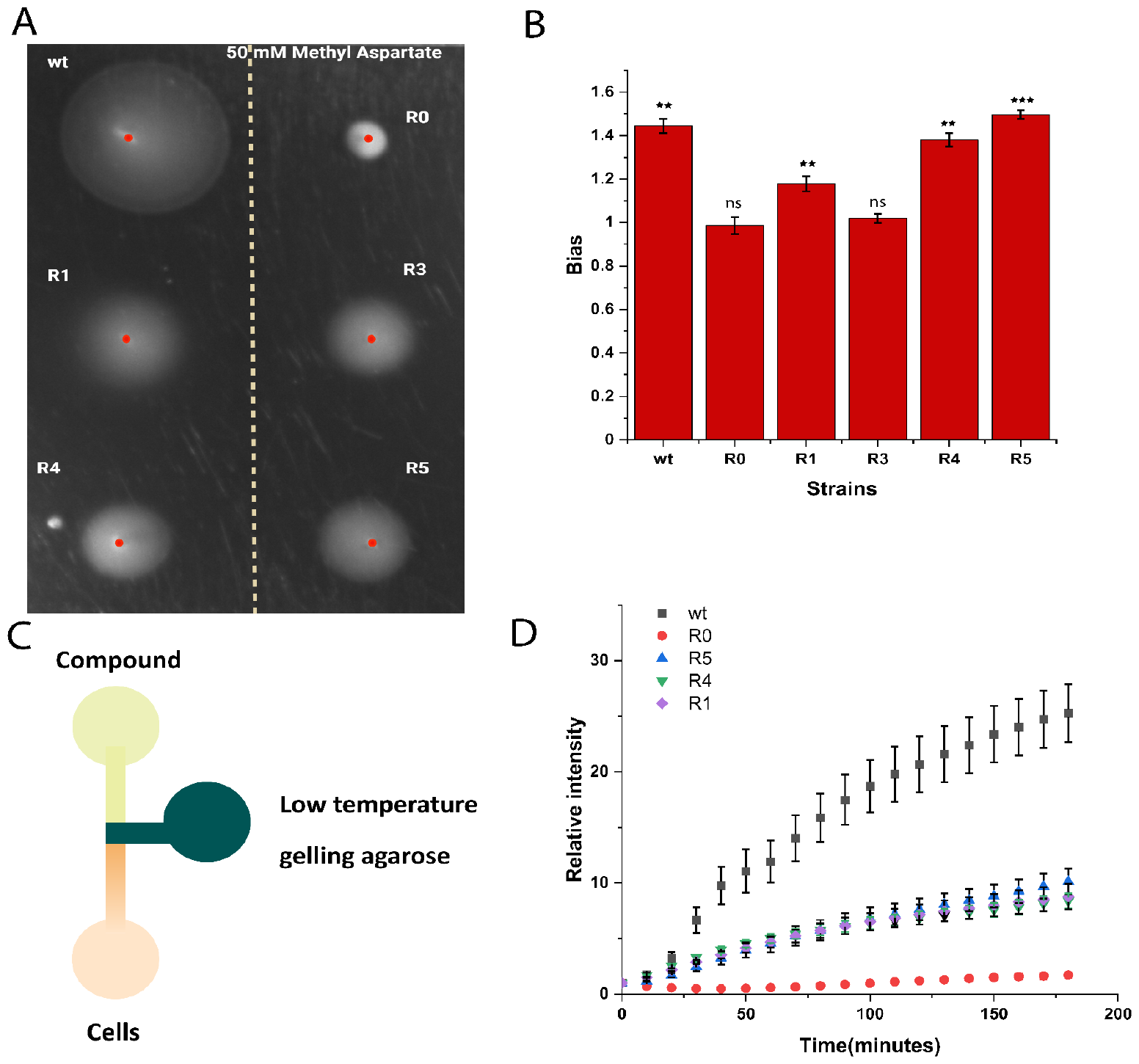
Biased movement of evolved Δ*cheR* strains towards sources of chemoattractant. **(A-B)** Indicated strains were tested for biased spreading on M9 minimal medium soft-agar (M9SA) plates with a pre-established gradient of α-methyl-D, L-aspartate (MeAsp), a non-metabolizable analog of aspartate, with 50 mM at the source (A). Spreading bias was measured in three independent replicates and quantified as the ratio between spreading up and spreading down the gradient. Error bars indicate standard errors. Significance analysis was done in comparison to the bias =1. *P* values were calculated using one-tailed t-test (ns, not significant; ^*^, *P <* 0.05; ^**^, *P <* 0.01; ^***^, *P <* 0.001). **(C-D)** Accumulation of fluorescently labels cells of indicated strains towards the source of tested compound (50 mM MeAsp) in the microfluidic device schematically represented in (C). Chemotactic accumulation is quantified as fluorescence intensity in the observation channel (depicted in orange in C, see also Figure S5) at indicated time points relative to the initial time point 0 (D).

To additionally confirm the ability of evolved strains to perform chemotaxis, we used a previously described microfluidic assay where an attractant gradient is generated in the liquid medium within the test channel (36-38) (Figure 4C). Consistent with these previous reports, chemotaxis was essential for efficient cell accumulation toward the source of MeAsp in this assay, since the wildtype strain accumulated rapidly whereas nearly no accumulation was observed for the fully motile but non-chemotactic Δ*cheR* (R0) strain (Figures 4D and Figure S5). In contrast, the tested evolved Δ*cheR* lines exhibited an intermediate but clear accumulation, suggesting that they can bias their swimming in a gradient of non-metabolizable attractant established in the liquid, albeit not as efficiently as the wildtype cells.

### Mechanism of the re-evolved chemotactic drift

In order to gain insight into the origin of this evolved chemotactic-like behavior, we analyzed the movement of individual swimming cells of the R1 line, as well as of the R0 cells, in a chemotactic chamber with or without a linear gradient of MeAsp (Figure 5A). In these experiments, tracks of motile cells of both strains showed no biased motion in the absence of a gradient (Figure 5B, C), as expected, and R0 cells also showed no bias in the gradients of MeAsp, established using either 100 µM or 1 mM at the source (Figure 5B). In contrast, cells of the R1 line showed a clearly pronounced chemotactic drift, particularly at the 1 mM gradient (Figure 5D).

**Figure 5.**
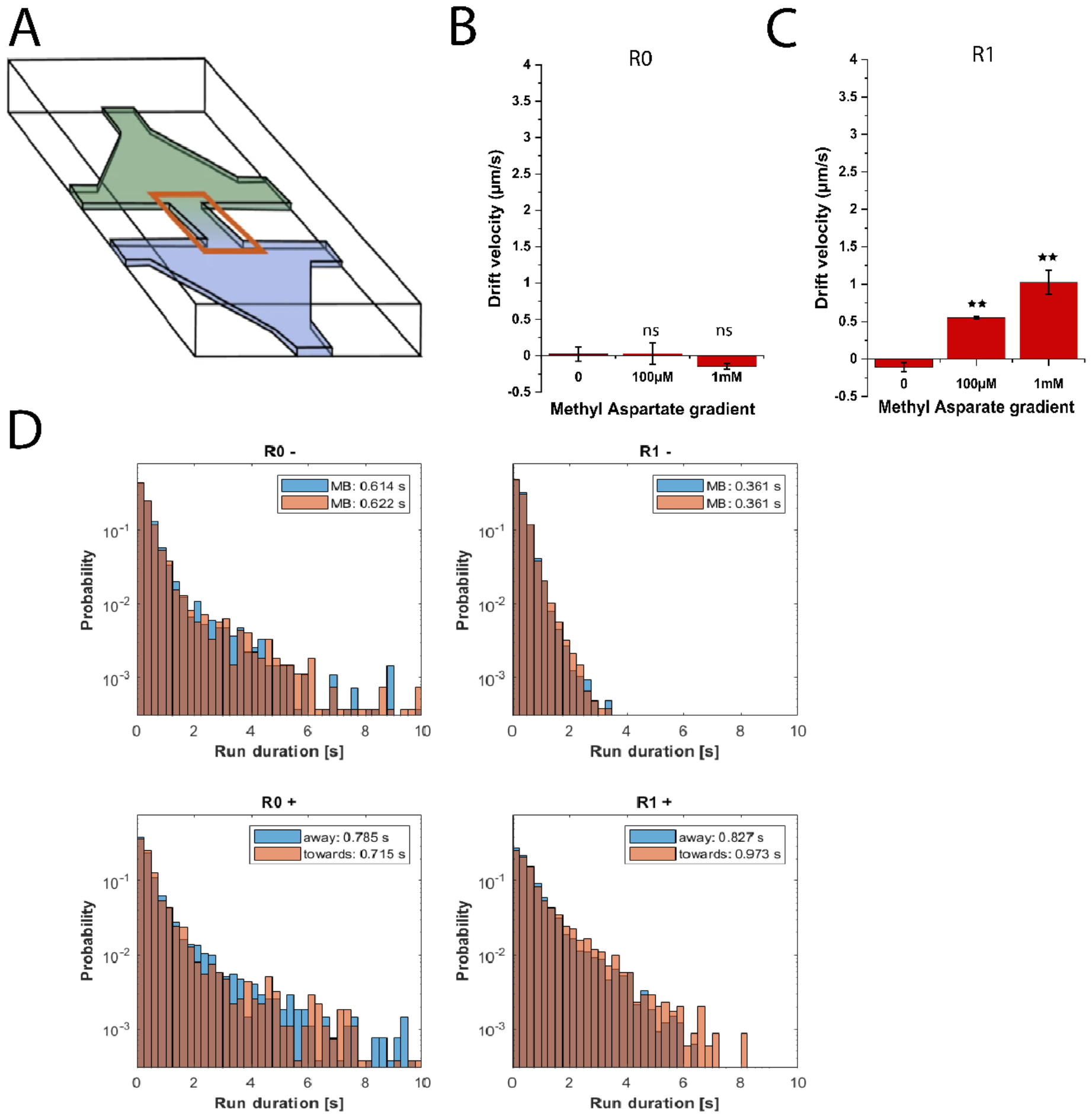
Chemotactic drift of R1 cells in a gradient of MeAsp. **(A)** Schematic of a chemotaxis microchamber made of poly-dimethylsiloxane (PDMS). Gradients in the observation channel were created by filling one chamber with motility buffer and the other with motility buffer containing either 100 µM or 1 mM MeAsp, or motility buffer as a negative control (0 gradient). Bacterial suspension was loaded in both chambers. **(B-C)** Drift velocity of Δ*cheR* (R0; B) and evolved R1 line (C) cells in these gradients, measured using differential dynamic microscopy (DDM). Significance analysis was done with respect to drift velocity values in absence of the gradient. *P* values were calculated using two tailed t-test (ns, not significant; ^*^, *P <* 0.05; ^**^, *P <* 0.01; ^***^, *P <* 0.001). **(D)** Distributions of cell run durations for indicated strains, either in absence of a gradient (-, top) or in presence of a gradient from zero to 1 mM MeAsp. Cells runs were measured using cell tracking and separated based on their direction, either towards or away from the source of chemoattractant. Mean duration of runs in either direction is indicated.

We further analyzed individual cell tracks for these strains (Figure 5E and Figure S6). In the absence of a gradient (-), both strains expectedly showed a similar distribution of their runs in both directions along the channel. The average run duration of the smooth-swimming Δ*cheR* cells was higher compared to the R1 cells, consistent with higher tumbling frequency of the R1 line (Figure 2A). In the presence of an attractant gradient, runs of the R0 cells became slightly longer on average, indicating a residual response of this strain to the presence of attractant. However, there was no difference between runs up and down the gradient, confirming that this strain is non-chemotactic. In contrast, runs of the R1 cells became strongly elongated both up and down the gradient of attractant, but showed a significantly stronger elongation up the gradient. This bias in the run length distribution is apparently sufficient to mediate the chemotactic drift in a gradient, even in the absence of the short-term adaptation.

Finally, we tested the pathway response of the evolved Δ*cheR* lines, using a previously described assay based on Förster (fluorescence) resonance energy transfer (FRET). This assay monitors the phosphorylation-dependent interaction between CheY fused to a yellow fluorescent protein (CheY-YFP) and its phosphatase CheZ fused to a cyan fluorescent protein (CheZ-CFP), as a readout of the pathway activity (39, 40). FRET measurements confirmed that the basal activity of the pathway increased in both lines tested, R1 and R4, compared to R0, allowing them to respond to MeAsp stimulation (Figure S7A). The sensitivity of the response was only moderately reduced compared to wildtype cells (Figure S7B,C). These evolved lines did not show any pronounced adaptation of pathway activity to sustained attractant stimulation, in contrast to wildtype cells, in which the recovery of pathway activity was very pronounced and followed by a characteristic overshoot upon the removal of attractant. Thus, despite their apparent ability to perform chemotactic-like navigation in a gradient, the evolved Δ*cheR* lines did not regain the ability to adapt to varying background stimulations on the timescale of our experiment. This inability of the R1 line to adapt to sustained stimulation was further confirmed by cell tracking in the presence of a uniform concentration of 500 µM MeAsp in the observation channel (Figure S6A).

## Discussion

Experimental evolution under defined laboratory selection has been used to provide important insights into the evolvability of individual proteins, simple traits, and regulatory networks in microorganisms (41-45), including the control of motility gene expression in bacteria (21-24). Here, we investigated the ability of experimental evolution to restore the function of the signaling network that controls bacterial chemotactic behavior in the absence of one of the essential components of the pathway. The *E. coli* chemotaxis pathway is one of the best understood signaling systems in biology, and all of its components are normally considered essential for its ability to mediate bacterial navigation in environmental gradients (1, 5, 11). This is consistent with the high conservation of all *E. coli* chemotaxis proteins across bacterial and archaeal phyla (46), with the sole exception of the phosphatase CheZ. However, the phosphatase activity itself is thought to be essential for the functionality of the pathway, as other bacteria possess alternative proteins that perform this function (9).

Partially consistent with this expected essentiality, we observed that the absence of the core signaling components, including CheA, CheY, CheW and chemoreceptors (MCPs), could not be restored by experimental evolution. Notably, CheA and CheY are likely to be the evolutionary oldest members of the pathway, directly related to the bacterial two-component systems (9). In contrast, cells deficient in auxiliary components, including the less conserved CheZ, but also the highly conserved but chemotaxis-specific adaptation enzymes CheR and CheB, recovered their ability to spread in soft agar, the assay typically used to assess the chemotactic ability of bacteria, albeit to varying degrees.

At the phenotypic level, an important but auxiliary factor for the improvement of spreading in soft agar was an increase in cell swimming velocity. A similar increase was already observed during the evolution of enhanced chemotaxis in wildtype cells with intact chemotaxis pathway (22). Consistent with this previous study, the increase in swimming could be explained by the upregulation of flagellar genes, either due to mutations in genes encoding components of the flagellar export apparatus or in regulatory genes controlling motility. However, introduction of the *fliI* mutation into the R1 line suggests that this increase in swimming alone cannot compensate for the defect in chemotaxis.

The more important phenotypic change was the restoration of tumbling behavior, with the optimal tumbling bias apparently close to that of the wildtype. This phenotypic adaptation could be attributed to the importance of intermediate tumbling frequencies for cells spreading in mesh-like soft agar pores, and such restoration was previously suggested to be sufficient to explain the spreading of Δ*cheR* revertant strains (14). At the molecular level, these changes in tumbling frequency could be explained by compensatory mutations in other chemotaxis genes that tune the activity of the pathway to an intermediate level. However, the observed mutations were clearly specific for restoring a particular deletion defect, strongly suggesting that their importance goes beyond simple modulation of pathway activity. For example, the tumbling phenotypes of Δ*cheZ* and Δ*cheB* mutants were compensated by different sets of mutations, and while the absence of *cheR* was commonly compensated by mutations in its counterpart *cheB*, the opposite was not true and no mutations in *cheR* could be observed in the evolved Δ*cheB* strain.

All this suggests that the observed evolutionary rewiring of the chemotaxis pathway does more than the simple restoration of the intermediate tumbling phenotype, but rather leads to a new strategy of simplified chemotactic behavior with a smaller set of pathway components. Indeed, evolved strains exhibited a clear bias in their behavior in chemoattractant gradients established in soft agar or in liquid. When the nature of this biased movement was examined for one of the evolved strains carrying the *cheR* deletion, we found that the behavior of this evolved strain differed from both the original Δ*cheR* and the wildtype strain. Whereas the wildtype strain showed strongly elongated runs up the attractant gradient, as previously reported (6), and also moderately shortened runs down the gradient, cells of both the original and the evolved Δ*cheR* strain extended their runs in both directions. This is consistent with the inability of these strains to rapidly adapt to the rapid changes in attractant concentration, which was confirmed for the evolved Δ*cheR* strain. However, although such adaptation is normally considered essential for chemotaxis, even in its absence the evolved Δ*cheR* strain showed greater elongation of runs up than down the gradient. This difference was apparently sufficient to produce a chemotactic drift in the gradient that was nearly half as efficient as that of wildtype cells. We further hypothesize that the residual activity of CheB retained in these evolved Δ*cheR* strains is required to gradually adjust the modification of their receptors, and thus the basal activity of the pathway, to the level of environmental stimulation through the synthesis of half-modified receptors and their gradual deamidation. Thus, by selectively modifying several chemotaxis proteins, the evolved *E. coli* strains developed a novel, adaptation-independent strategy of chemotaxis.

## Materials and Methods

### Strains and plasmids

*Escherichia coli* strains listed in Table S4 were grown at 37°C in Luria broth (LB) with 200 rpm shaking or on LB containing 1.5% (w/v) agar (LBA). For spreading ring measurements, *E. coli* strains were grown at 30°C in Tryptone broth (TB) containing 0.27% (w/v) agar (Tryptone Broth Soft Agar-TBSA).

### Experimental evolution of the chemotaxis mutants

The chemotaxis mutants (Δ*cheA*, Δ*cheB*, Δ*cheR*, Δ*cheW*, Δ*cheY*, Δ*cheZ*, and receptor-less strain) were evolved through 30 passages of sub-culturing in TBSA plates. 2µl cells from the outer edge of the spread ring after overnight growth at 30°C (on the TBSA plate) were inoculated onto a fresh TBSA plate. This process was repeated daily for 30 days, with an additional sample taken for glycerol stocks.

### Genome editing

The recipient strain containing pKD46 was transformed with pKD45 with linear DNA of the targeted product using electroporation. Transformed cells were grown in LB + 0.2% Arabinose for 4-5 hrs at 30°C. These cells were plated on Rhamnose plates for selection. Rhamnose-resistant colonies were selected and sequenced to confirm.

### Re-sequencing of bacterial genomes

The genomic DNAs of the evolved bacterial population were isolated using the Qiagen DNeasy Blood and Tissue kit following the manufacturer’s instructions. Libraries were constructed using Nextera XT Index Kit (24 indexes, 96 samples) Illumina FC-131-1001. Sequencing was done using the Illumina HiSeq Rapid Run (2 × 150 bp paired-end run). The genomes were reassembled with the DNASTAR Seqman NGen 12 software and BRESEQ pipeline using the *E*.*coli* RP437 genome as the template. Genes containing point mutations were amplified using PCR. This PCR fragment was subsequently cleaned and sent for Sanger Sequencing to Microsynth Seqlab GmbH.

### Gradient plate assay

Minimal A agar (0.25% agar, 10 mM KH_2_PO_4_/K_2_HPO_4_, 8 mM (NH_4_)_2_SO_4_, 2 mM citrate, 1 mM MgSO_4_, 0.1 mg mL^−1^ of thiamine-HCl, 1 mM glycerol, and 40 μg mL^−1^ of a mixture of threonine, methionine, leucine, and histidine) supplemented with antibiotics and inducers was used for the agar plate assay. Chemical solutions (50mM serine and methyl aspartate) were applied to the centerline of the plate and incubated at 4 °C for 12–16 h to generate a chemical gradient before cell inoculation. Overnight cultures of different evolved strains were washed twice with tethering buffer and applied to the plate at a distance of 1.5 cm from the line where the chemical was inoculated. Plates were incubated at 30 °C for 24–48 h. Spreading bias was quantified by measuring the spreading of the bacterial population up or down the chemical gradient using ImageJ analysis.

### Microfluidics chemotaxis assay

Cells were harvested by centrifugation at 4000 rpm for 5 min and washed twice with the tethering buffer. Methyl aspartate dissolved in the tethering buffer (50mM) was adjusted to pH 7.0. The responses of *E. coli* cells to concentration gradient were measured using a microfluidic device. In summary, *E. coli* cells were added at the sink side pore of the device to a final OD_600_ of 1.2–2 and equilibrated for 20 min in the observation channel. Methyl Aspartate solution was added at the source side pore and allowed to diffuse into the observation channel for an indicated time to establish a concentration gradient. Fluorescence microscopy on a Nikon Ti-E microscope system with a 20× objective lens was used to detect the fluorescence intensity of cells in the observation channel. The cellular response was characterized by the fluorescence intensity in the observation channel’s analysis region (300 μm × 200 μm). Data were analyzed using ImageJ (Wayne Rasband, National Institutes of Health, USA).

### FRET assay

Bacterial cells were transformed with pVS88 plasmid for FRET assay experiment and were grown in TB (1% tryptone, 0.5% NaCl) supplemented with antibiotics (100 μg mL^−1^ of ampicillin; 17 μg mL^−1^ of chloramphenicol) and appropriate inducers at 34 °C and 275 rpm. Cells were harvested at OD_600_ of 0.6 and washed twice with the tethering buffer (10 mM KH_2_PO_4_/K_2_HPO_4_, 0.1 mM EDTA, 1 μM methionine, 10 mM sodium lactate, pH 7.0). FRET measurements were performed on an upright fluorescence microscope (Zeiss Axio Imager.Z1). Strains were stimulated with compounds of interest. The fluorescence signals were recorded, analyzed as described previously (5), and plotted using KaleidaGraph (Synergy Software). Data were fit to a Hill model. For the repellent response, *Y* = *A* × *L*^H^/(*L*^H^ + *K*^H^), whereas for the attractant response, *Y* = *A* × (1 – *L*^H^/(*L*^H^ + *K*^H^)). In the model, *Y* is the initial FRET response, *L* is the ligand concentration, *A* is the amplitude (for the saturated response, *A* is fixed to be 1), *H* is the Hill coefficient, and *K* is the EC_50_.

### Promoter Activity Analyses

For promoter activity assays, *E. coli* strains transformed with reporter plasmid pAM109 and grown in TB supplemented with kanamycin in 96-well plates at 30°C in a rotary shaker at 180 rpm. Cell fluorescence was measured using flow cytometry on BD LSR Fortessa SORP cell analyzer (BD Biosciences).

### Motility analyses

For motility analysis, *E. coli* cells were grown in 10 mL TB medium at 34°C in a rotary shaker at 180 rpm. Cells (1 mL) at mid-log phase (OD600 [optical density at 600 nm] = 0.6) were collected and re-suspended in 1 mL tethering buffer (10 mM K_2_HPO_4_, 10 mM KH_2_PO_4_, 100 mM EDTA, 1 mM L-methionine, 10 mM lactic acid [pH 7.0]). Swimming velocity and tumbling rate were measured by cell tracking in a glass chamber using phase-contrast microscopy (Nikon TI Eclipse, 103 objective, NA = 0.3, CMOS camera EoSens 4CXP). All data were analyzed using ImageJ (https://imagej.nih.gov/ij/) with custom-written plugins for swimming velocity, drift velocity, and tumbling rate analysis (53).

## Acknowledgments

We thank Ferencz Paldy and Lars Velten for their help with preliminary experiments for this study. This work was supported by the Max Planck Society.

