## Supplementary Materials for "Experimental evolution of a reduced bacterial chemotaxis network"

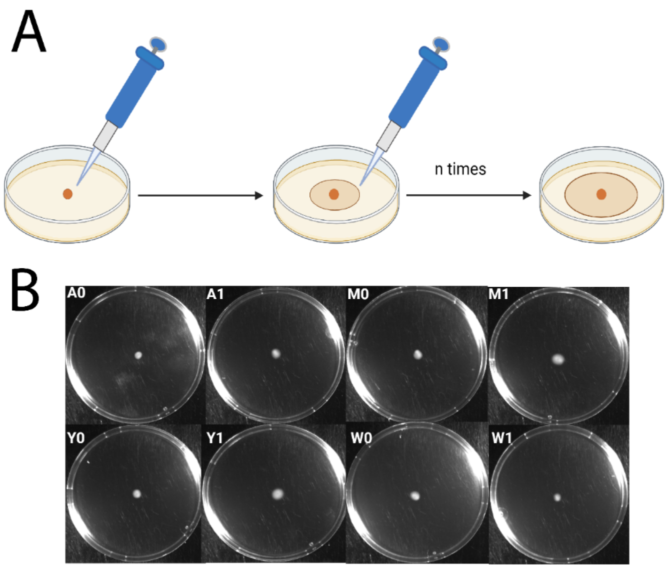


**Figure S1: Methodology of the evolution experiment and spreading of core chemotaxis gene deletion strains.** **(A)** Schematic of the evolution experiment, where the fastest spreading cells from the edge of the spreading ring were collected and inoculated in the middle of a fresh TBSA plate, with the  for 30 days. **(B)** Spreading of Δ*cheA* (A), Δ*cheY* (Y), Δ*mcp* (M) or Δ*cheW* (W) deletion strains, either before (denoted as “0”) or after evolution for 30 days in soft agar, with an example of one evolved line (denoted as “1”) shown for each strain.

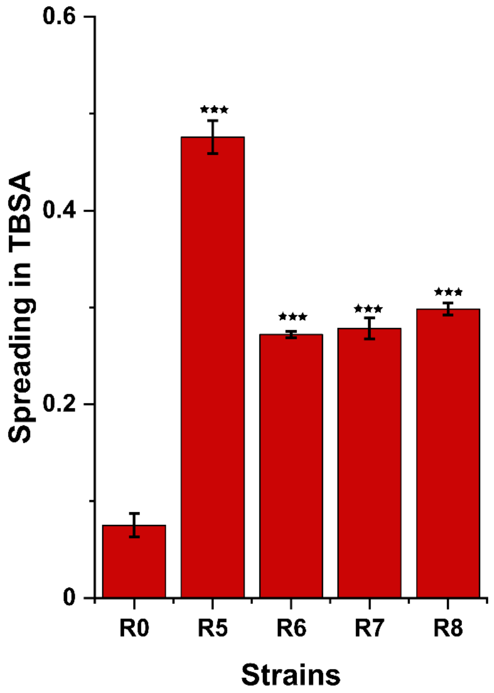


**Figure S2: Size of the spreading colonies for additional evolved Δ*cheR* lines.** Spreading was measured for indicated R lines in three independent replicates after ~16 h incubation in TBSA and normalized to spreading of the wildtype (wt). Error bars indicate standard errors. *P* values were calculated with respect to the non-evolved Δ*cheR* strain (R0) using a two-tailed t-test (ns, not significant; *, $P < 0.05$; **, $P < 0.01$; ***, $P < 0.001$).

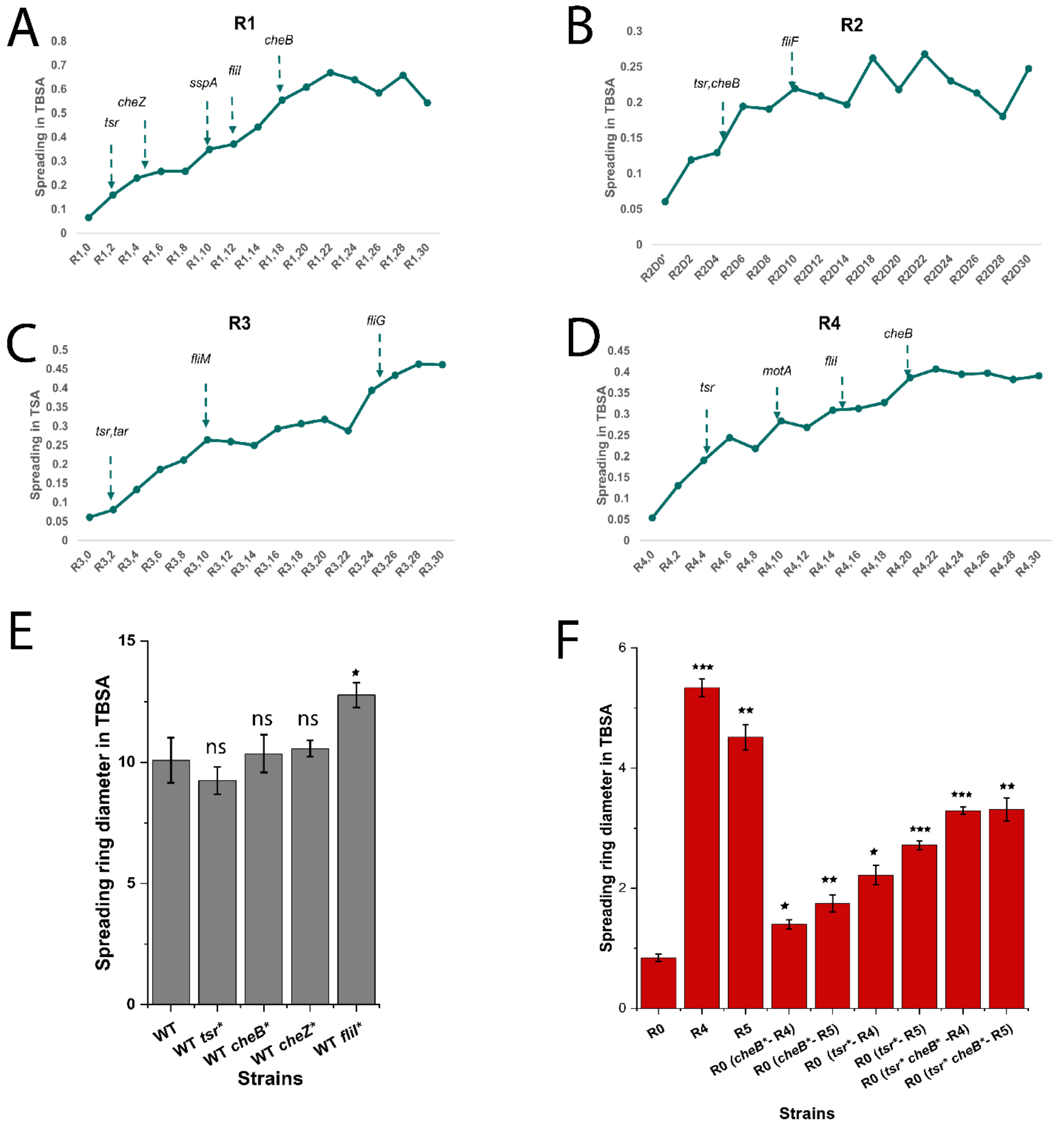


**Figure S3: Order of appearance and effects of point mutations in Δ*cheR* lines. (A-D)** Spreading of R1 (A), R2 (B), R3 (C), and R4 (D) lines in TBSA over the time course of experimental evolution (in days), with the first day when respective mutation could be identified by Sanger sequencing being highlighted. Spreading values are from a single experiment and normalized to spreading of wt. **(E)** Effects of individual R1 mutations on spreading when introduced in wt background. Values were measured in three independent replicates and normalized to spreading of wt. Error bars indicate standard errors. Significance analysis was done in comparison to wt. *P* values were calculated using two tailed t-test (ns, not significant; *, $P < 0.05$; **, $P < 0.01$; ***, $P < 0.001$).

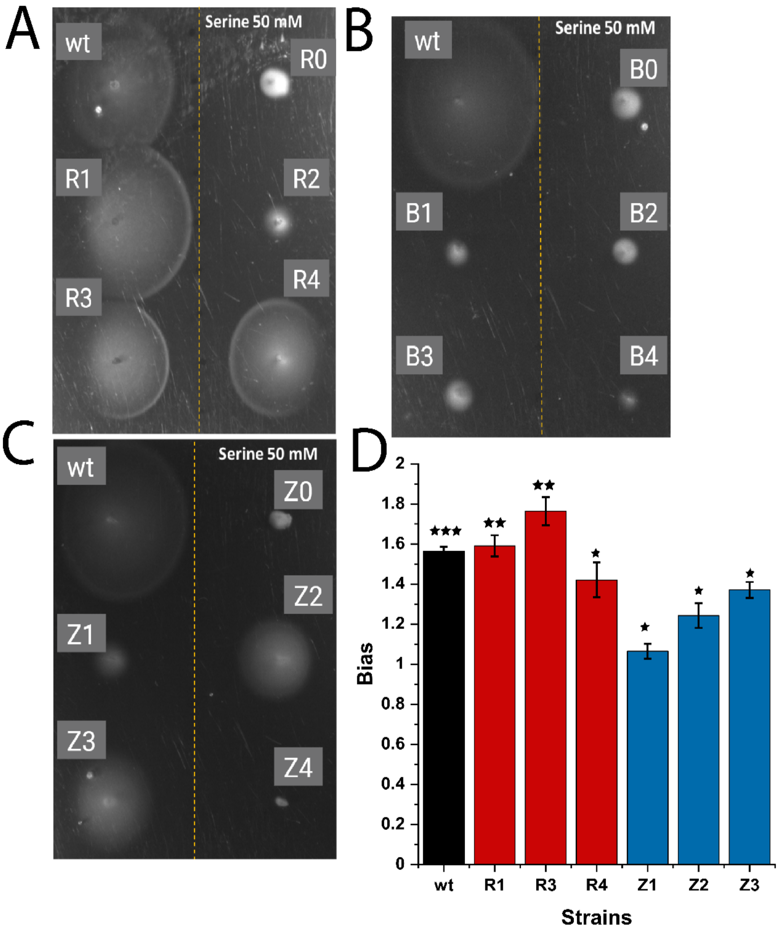


**Figure S4: Biased movement of evolved strains towards serine. (A-C)** Indicated strains were tested for biased spreading on M9 minimal medium soft-agar (M9SA) plates with a pre-established gradient of serine (50 mM at the source). **(D)** Spreading bias was measured in three independent replicates and quantified as the ratio between spreading up and spreading down the gradient. Error bars indicate standard errors. Significance analysis was done in comparison to the bias =1. *P* values were calculated using one-tailed t-test (ns, not significant; *, $P < 0.05$; **, $P < 0.01$; ***, $P < 0.001$).

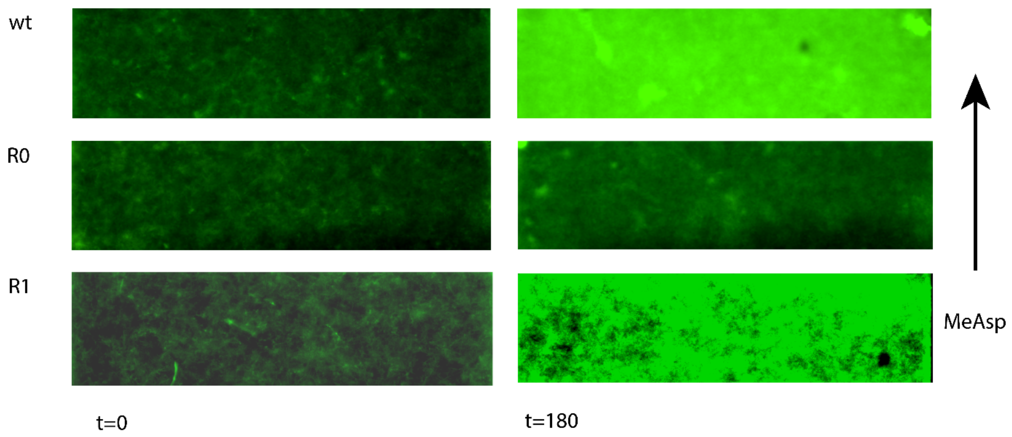


**Figure S5: Images of the observation channel of the microfluidic device used to quantify chemotaxis in Figure 4C and D.** Images for indicated strains and times are shown. Arrow points to the direction in which source of chemoattractant (MeAsp) is located.

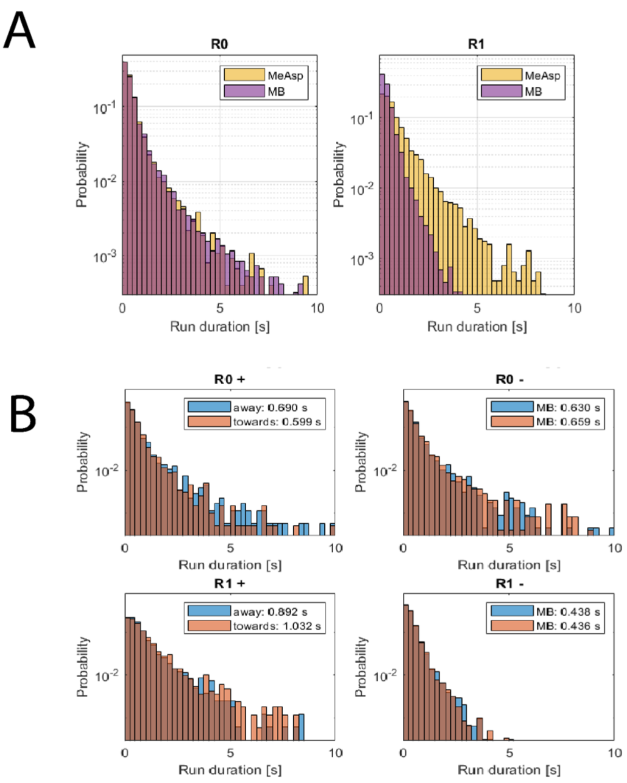


**Figure S6: Run duration in presence of steady stimulation.** Distributions of cell run durations for indicated strains. **(A)** Cells were adapted either in motility buffer (MB) or in presence of spatially uniform stimulation with 500 µM MeAsp. **(B)** Cells were exposed to a gradient of MeAsp from zero to 1 mM. Cells runs were measured using cell tracking and separated based on their direction, either towards or away from the source of chemoattractant. Mean duration of runs in either direction is indicated.

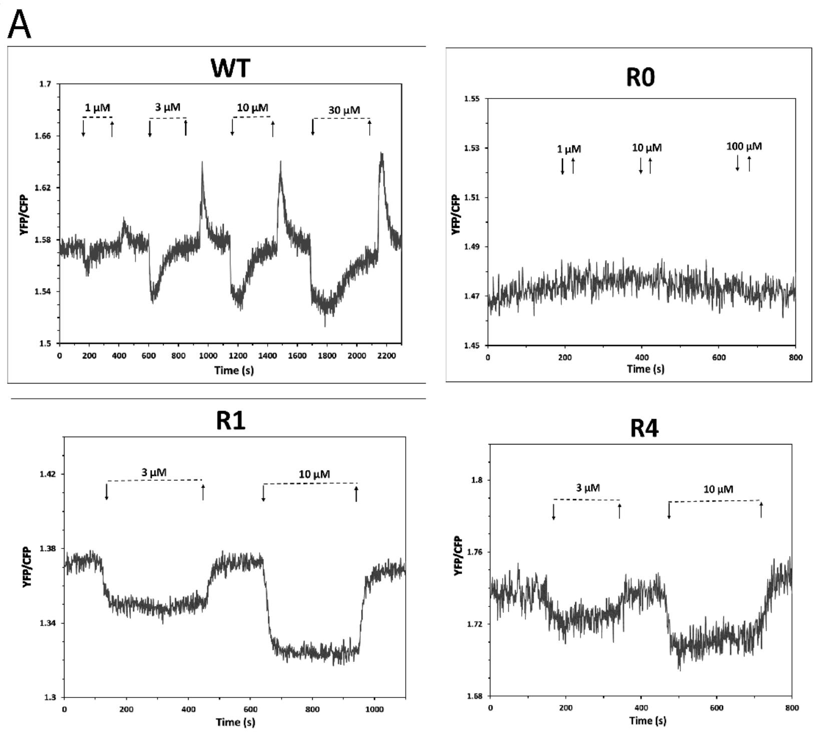


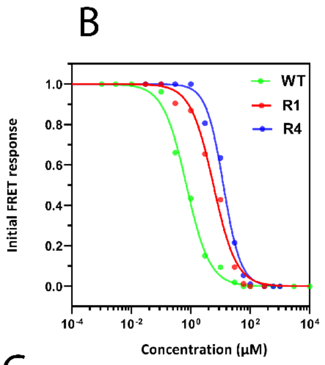


C

| Strain | EC50 |
|---|---|
| WT | 0.71 ± 0.05 μM |
| R1 | 5.79 ± 1.34 μM |
| R4 | 12.57 ± 1.97 μM |

**Figure S7: Pathway response in wildtype and evolved Δ*cheR* strains to MeAsp measured by FRET. (A)** FRET measurement of responses to MeAsp for wt and indicated R strains, with the ratio between signals in yellow and cyan fluorescence channels being proportional to pathway activity. Indicated concentrations of MeAsp were added (down arrow) and subsequently removed (up arrow). No response was observed for R0 strain, consistent with its very low basal pathway activity. **(B,C)** Dose-response measurements for MeAsp stimulation of indicated strains (B) and corresponding values of the half-inhibitory concentration (EC50) resulting from these measurements (C).

**Table S1: Whole genome sequencing revealed multiple mutations in the evolved Δ*cheR* lines.**

| gene | annotation | R1 | R2 | R3 | R4 | R5 | R6 | R7 | R8 |
|---|---|---|---|---|---|---|---|---|---|
| *tsr* | N54D | | | ■ | | | | | |
| *tsr* | V199M | | | | | | | | ■ |
| *tsr* | M249T | | ■ | | | | | | |
| *tsr* | L263M | | | | ■ | | | | |
| *tsr* | T305M | ■ | | | | | | | |
| *tsr* | G426E | | | | | | ■ | | |
| *tsr* | T441K | | | | | ■ | | | |
| *tar* | A166P | | | ■ | | | | | |
| *tar* | V433I | | | | | | | ■ | |
| *cheZ* | Q64L | | | | | | ■ | | |
| *cheZ* | Q204L | ■ | | | | | | | |
| *cheB* | S8P | | | | | | ■ | | |
| *cheB* | L71F | | | | ■ | | | | |
| *cheB* | R75C | ■ | | | | | | | |
| *cheB* | A130V | | | | | | | | ■ |
| *cheB* | R257G | | ■ | | | | | | |
| *cheB* | G313C | | | | | ■ | | | |
| *motA* | L2F | | | | ■ | | | | |
| *fliF* | G520R | | ■ | | | | | | |
| *fliG* | E114Q | | | ■ | | | | | |
| *fliG* | G165S | | | | | | | | ■ |
| *fliG* | G165A | | | | | ■ | | | |
| *fliI* | T42I | | | | | | | | ■ |
| *fliI* | M80I | | | | ■ | | | | |
| *fliI* | M178I | ■ | | | | | | | |
| *fliI* | R197H | | | | | | | ■ | |
| *fliM* | V98L | | | | | | | ■ | |
| *fliM* | P195L | | | ■ | | | | | |
| *rsfS / cobC* | IS1 | | | | ■ | | | | |
| *lysO* | P241L | | | ■ | | | | | |
| *ycjM* | IS1 | | ■ | | | | | | |
| *ddpB* | I6T | ■ | | | | | | | |
| *rsmI* | P101L | | ■ | | | | | | |
| *[nanA]–[sspA]* | Δ4,084 bp | | | ■ | | | | | |
| *sspA* | 14 bp insertion | ■ | | | | | | | |
| *atpI / rsmG* | IS1 | | ■ | | | | | | |
| *fabR* | IS1 | | | | ■ | | | | |
| *perR* | IS1 | | | | | | | | |
| *[gltI]* | IS5 | ■ | | | | | | | |
| *mngB / cydA* | IS5 | | | | | ■ | | | |
| *opgG* | T405K | | | | | | | | ■ |
| *purR* | Δ20 bp | | | | | | | | |
| *eda* | W4C | | | | | | ■ | | |
| *edd* | 5 bp insertion | | | | | | | | ■ |
| *edd* | S16* | | | | | ■ | | | |
| *[rbn]* | IS1 | | | | | | | | ■ |
| *ppk* | IS1 | | | | | | | | ■ |
| *parC* | A414S | | | | | | ■ | | |
| *sspA* | IS1 | | | | | ■ | | | |
| *gntT* | IS1 | | | | | | | | |
| *rplI* | Q133* | | | | | | ■ | | |

**Table S2: Whole genome sequencing revealed multiple mutations in the evolved Δ*cheB* lines.**

| gene | annotation | B1 | B2 | B3 | B4 |
|---|---|---|---|---|---|
| *tsr* | A94T | | | | ■ |
| *tsr* | S437R | ■ | | | |
| *tap* | S428L | | | ■ | |
| *tar* | A411T | | ■ | | |
| *tar* | A496-A498 deletion | | | | ■ |
| *cheW* | D139(ALGD insertion) | | ■ | | |
| *cheA* | E319A | ■ | | | |
| *fliN* | A115S | | | ■ | |
| *rpsA* | R86H | | ■ | | |
| *yciW* | E217K | | | ■ | |
| *ydcO* | IS1 | | | ■ | |
| *ddpB* | I6T | | | | |
| *fadD* | IS5 | ■ | | | |
| *fadD* | 76 bp deletion | | | | ■ |
| *atpD* | L163R | | | | |
| *atpA* | D289H | ■ | | | |
| *atpH* | G84H | | | | ■ |
| *atpI* / *rsmG* | IS5 | | | ■ | |
| *atpI* / *rsmG* | IS1 | | ■ | | |

**Table S3: Whole genome sequencing revealed multiple mutations in the evolved Δ*cheZ* lines.**

| gene | annotation | Z1 | Z2 | Z3 | Z4 |
|---|---|---|---|---|---|
| *tar* | S31 frameshift | | | ■ | ■ |
| *tar* | Q155* | ■ | ■ | | |
| *cheA* | L92W | ■ | | | |
| *cheA* | M98V | | | | ■ |
| *cheA* | P457L | | | ■ | |
| *cheA* | D476E | | ■ | | |
| *rrsH* | frameshift | ■ | | | |
| *clpX* | IS1 | ■ | ■ | | |
| *rhsC* | 2 bp deletion | | | ■ | |
| *opgH* | V463G | | | ■ | |
| *topA* | L781Q | | | ■ | |
| *bamD* | Y205H | | | ■ | |
| *rpoD* | 27 bp insertion | | | ■ | |
| *atpD* | L163R | ■ | | | |
| *atpI* / *rsmG* | IS5 | | | | ■ |

**Table S4: Strains and plasmids used in this study.**

| Name | Strain genetic background | Source |
|---|---|---|
| **Strains** | | |
| RP437 (WT) | *Escherichia coli* RP437 (wild type for chemotaxis) | (1) |
| VS126(R0) | RP437 Δ*cheR* | (2) |
| RP4972(B0) | RP437 Δ*cheB* | (1) |
| VS161(Z0) | RP437 Δ*cheZ* | (2) |
| VS166(A0) | RP437 Δ*cheA* | (3) |
| VS100(Y0) | RP437 Δ*cheY* | (4) |
| VS289(W0) | RP437 Δ*cheW* | (1) |
| UU1250(M0) | RP437 Δ*tar* Δ*tap* Δ*tsr* Δ*aer* Δ*trg* | (5) |
| VS1936 | R1 Δ*cheB* | This study |
| VS1956(R0 *cheB**) | RP437 Δ*cheR cheB** (R75C) | This study |
| VS1957(WT *cheB**) | RP437 *cheB** (R75C) | This study |
| VS1958(WT *cheZ**) | RP437 *cheZ** (Q204L) | This study |
| VS1959(R0 *cheZ**) | RP437 Δ*cheR cheZ** (Q204L) | This study |
| VS1960(R0 *tsr*cheZ**) | RP437 Δ*cheR tsr** (T305M) *cheZ** (Q204L) | This study |
| VS1961(R0 *cheB*cheZ**) | RP437 ΔcheR *cheB** (R75C) *cheZ** (Q204L) | This study |
| VS1962(R0 *cheB*tsr**-R4) | RP437 Δ*cheR cheB**( L71F) *tsr** (L263M) | This study |
| VS1963(R0 *cheB*tsr**-R5) | RP437 Δ*cheR cheB** (G313C) *tsr** (T441K) | This study |
| VS1964(WT *fliI**) | RP437 *fliI** (M178I) | This study |
| VS1965(R0 *fliI**) | RP437 Δ*cheR fliI** (M178I) | This study |
| VS1966(R0 *tsr*cheZ*fliI*)* | RP437 Δ*cheR tsr**( T305M) *cheZ** (Q204L) *fliI** (M178I) | This study |
| VS1968(R0 *tsr**-R4) | RP437 Δ*cheR tsr** (L263M) | This study |
| VS1970(R0 *tsr**-R5) | RP437 Δ*cheR tsr** (T441K) | This study |
| VS1972(R0 *cheB**-R4) | RP437 Δ*cheR cheB** (L71F) | This study |
| VS1974(R0 *cheB**-R5) | RP437 Δ*cheR cheB** (G313C) | This study |
| VS1975(R0 *tsr*cheB*)* | RP437 Δ*cheR tsr** (T305M) *cheB**( R75C) | This study |
| VS1976(R0 *tsr*cheB*cheZ*)* | RP437 Δ*cheR tsr** (T305M) *cheZ** (Q204L) *cheB** (R75C) | This study |
| VS1977(R0 *tsr*cheB*cheZ*fliI**) | RP437 Δ*cheR tsr** (T305M) *cheZ** (Q204L) *fliI** (M178I) *cheB** (R75C) | This study |
| **Plasmids** | | |
| pVS88 | CheY-EYFP / CheZ-ECFP expression plasmid used for FRET assays | (6) |
| pAM109 | GFP reporter for *fliC* promoter was constructed based on pUA66 | (7) |
| pKD46 | Used for making chromosomal deletions of genes with FRT sites | (8) |
| pKD45 | Used for SNP introduction; counterselection with *ccdB* gene under a rhamnose-inducible promoter while introducing | J.S.Parkinson, personal gift and (8) |
